# Identifying drivers of parallel evolution: A regression model approach

**DOI:** 10.1101/118695

**Authors:** Susan F. Bailey, Qianyun Guo, Thomas Bataillon

**Affiliations:** Bioinformatics Research Centre, Aarhus University, C.F. Møllers Allé 8, DK-8000 Aarhus C, Denmark.; Department of Biology, Clarkson University, PO Box 5805, Potsdam, NY 13699-5805

**Keywords:** parallel evolution, experimental evolution, Poisson regression, negative binomial regression

## Abstract

This preprint has been reviewed and recommended by Peer Community In Evolutionary Biology (http://dx.doi.org/10.24072/pci.evolbiol.100045). Parallel evolution, defined as identical changes arising in independent populations, is often attributed to similar selective pressures favoring the fixation of identical genetic changes. However, some level of parallel evolution is also expected if mutation rates are heterogeneous across regions of the genome. Theory suggests that mutation and selection can have equal impacts on patterns of parallel evolution, however empirical studies have yet to jointly quantify the importance of these two processes. Here, we introduce several statistical models to examine the contributions of mutation and selection heterogeneity to shaping parallel evolutionary changes at the gene-level. Using this framework we analyze published data from forty experimentally evolved *Saccharomyces cerevisiae* populations. We can partition the effects of a number of genomic variables into those affecting patterns of parallel evolution via effects on the rate of arising mutations, and those affecting the retention versus loss of the arising mutations (i.e. selection). Our results suggest that gene-to-gene heterogeneity in both mutation and selection, associated with gene length, recombination rate, and number of protein domains drive parallel evolution at both synonymous and nonsynonymous sites. While there are still a number of parallel changes that are not well described, we show that allowing for heterogeneous rates of mutation and selection can provide improved predictions of the prevalence and degree of parallel evolution.

**Data archival location:** Dryad, doi to be included later

## Introduction

Documenting patterns of parallel evolution during the adaptive divergence of populations or during repeated bouts of adaptation in populations maintained in the lab is becoming increasingly feasible. Beyond the fascination for the pattern of repeatable evolution, an outstanding open question is to understand which underlying processes are driving the pattern of molecular evolution during adaptation. Theory makes clear cut predictions: in the absence of selective interference between beneficial mutations (the so called strong selection weak mutation, or SSWM, domain), heterogeneity in mutation rates and selection coefficients between loci are expected to have equal influence on patterns of parallel evolution (Chevin et al., 2010; Lenormand et al., 2016). So far very few empirical studies have attempted to jointly quantify the relative importance of these two processes in shaping patterns of parallel evolution in genetic data. One study has explored this indirectly by quantifying the contribution of these two processes in shaping the parallel evolution of heritable traits that are assumed to be associated with parallel genetic changes (Streisfeld and Rausher, 2011). Recent work by Bailey et al., 2017 outlines an approach for quantifying the effects of mutation and selection heterogeneity in driving parallel evolution in experimental evolution data, but this alternate approach can not identify potential genomic drivers of that heterogeneity, as we do here. Other previous studies looking explicitly at parallel genetic changes have focused on the impacts of either selection or mutation separately.

Parallel evolution is an identical change in independently evolving lineages, and the similar process, convergent evolution, occurs when different ancestral states change to the same descendant state in independently evolving lineages (Zhang and Kumar, 1997). These kinds of evolutionary changes are studied across many different levels of biological organization from nucleotides to genes to pathways and more. In this study, we focus on parallel evolution at the level of the gene.

Parallel, along with convergent, evolution has previously been considered strong evidence of selection. A number of studies have examined gene-level mutation counts, looking for levels of parallel evolution that exceed what one would expect in the absence of selection (Caballero et al., 2015; Marvig et al., 2015; Woods et al., 2006), according to some null model, with an aim to identify genes that are under selection. For example, (Caballero et al., 2015) calculated the probability of instances of gene-level parallel evolution in whole genome sequences of *Pseudomonas aeruginosa* repeatedly sampled over the course of a year from the sputum of a cystic fibrosis patient assuming uniform re-sampling of ~150 mutation events across the approximately 6000 genes in the genome. The authors were able to identify 19 different genes for which there was significant deviation from their null model, and that pattern was interpreted as evidence for selection acting on these genes. However this study and other similar approaches do not account for the possibility of heterogeneity in mutation rate from gene-to-gene, a process that can generate false positives when using “abnormal” levels of parallel evolution as a means to detect selected genes.

Others have compared instances of parallel and convergent evolution across species (see Christin et al., 2010 for a review and examples). These studies also aim to identify genes under selection by searching for genes that exhibit a higher than expected number of instances of parallel evolution according to a specified null model for evolution. Many cross-species comparative studies report instances of parallel molecular evolution and readily interpret these as being driven by positive selection (e.g. Castoe et al., 2009; Feldman et al., 2012; Jost et al., 2008; Liu et al., 2014). However Zou and Zhang, 2015 show that in this type of analysis the choice of null model is crucial and suggest that many previously reported instances of parallel evolution driven by selection could in fact have resulted simply from mutation biases and mutational heterogeneity in the absence of selection.

In contrast to studies aimed at identifying selection, other work has focused on examining how heterogeneity in mutation rate can effect the distribution of mutations across a genome, and so the probability of parallel evolution. These studies focus exclusively on either those mutations that are assumed to be to a first approximation neutral (e.g. synonymous mutations, Maddamsetti et al., 2015) or mutations arising in the course of experiments where selection is minimal (e.g. mutations arising in a mutation accumulation experiment, Ness et al., 2015). On the whole, these studies suggest substantial gene-to-gene heterogeneity in mutation rate and this can arguably also generate differences in the distribution of mutations across the genome (although studies differ in the factors identified that drive that heterogeneity). However, it is not clear what the relative contribution of mutation rate heterogeneity is when the mutations of interest also have the potential to be under varying degrees of selection.

In this study we aim to explore the effects of both mutation and selection in generating the mutations that are observed across the genome. By identifying and quantifying the processes that give rise to mutations and how those vary from gene-to-gene, we can then begin to understand and predict patterns of parallel evolution at the gene-level. To do this we propose a framework that explicitly considers drivers of both selection and mutational heterogeneity. Using both Poisson and negative binomial regression models, we analyze gene-level mutation count data obtained from whole genome sequencing of a large set of yeast (*Saccharomyces cerevisiae*) experimental populations that were adapted in parallel to identical environmental conditions in the lab (Lang et al., 2013). We find that the best predictor of parallel mutations at the gene-level is simply the length of the gene, and along with this, a few other genomic covariates – namely the number of protein domains and the rate of recombination, and so it is variation in these variables that drives patterns of parallel evolution in this system.

## Models for identifying processes underlying parallel evolution

We are interested in quantifying heterogeneity in mutation rate and selection, and how these in turn are driving patterns of parallel evolution, and identifying genomic variables that predict how the processes of mutation and selection vary from gene-to-gene. To accomplish this, we need a framework that can explicitly separate the effects of variation in mutation rate and variation in selection. We do this by examining separately the observed synonymous and nonsynonymous mutations, making the assumption (which we then explore) that gene-to-gene variation in the rate at which synonymous mutations rise to observable frequencies is driven solely by variation in the mutation rate per gene, while gene-to-gene variation in the rate at which nonsynonymous mutations have arisen may be driven by heterogeneity in both mutation and selection processes. We describe the number of mutations observed in gene *i* during the course of an experiment, as 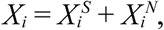 where 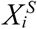 and 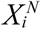 denote the synonymous and nonsynonymous mutation counts respectively. We assume these mutations are Poisson distributed with rates 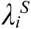 and 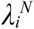 respectively. For synonymous mutations, this Poisson rate can be modeled as

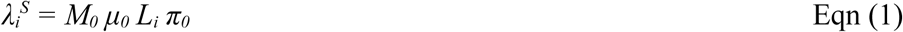

Here, *M*_*0*_ is a parameter that absorbs both time and population size at which the evolution occurred and that is constant across the genome, *μ*_*0*_ is the per-nucleotide mutation rate that we assume (and check) is constant across the genome, *L*_*i*_ is the length of gene *i* in nucleotides, and *π*_*0*_ is the probability of a synonymous mutation rising to an observable frequency in the population (we assume that synonymous mutations are selectively neutral and so this probability is assumed to be constant across the genome). For nonsynonymous mutations,

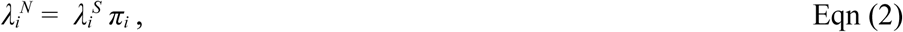

where *π*_*i*_ is a scalar that incorporates the effects of selection on the rate of fixation of non-synonymous mutations arising in gene *i*. Specifically, *π*_*i*_, is a function of the mean selection coefficient of gene *i, s*_*i*_, and under strong-selection-weak-mutation (SSWM) conditions, *π*_*i*_ ∝ *s*_*i*_ (Gillespie, 1984). We assume that the mean selection coefficient for non-synonymous mutations in a given gene can range from deleterious, to neutral, to beneficial. The type of data used and the underlying assumptions are summarized in Fig. 1.

**Figure 1:**
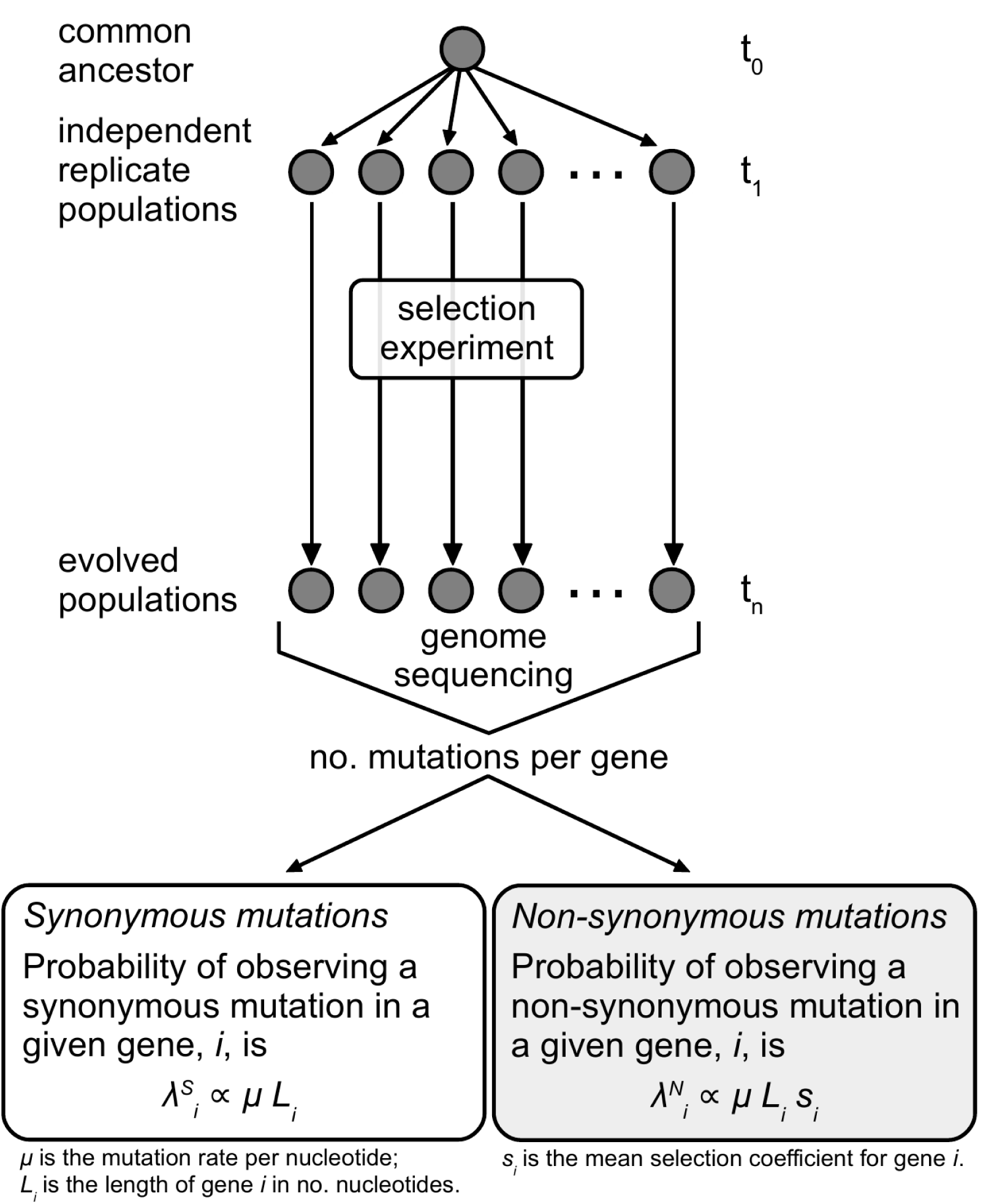
Schematic showing how the mutation counts data are generated and general assumptions underlying these data.

Given these underlying assumptions about the processes giving rise to observable mutations in the experimental sequence data, we can then use Poisson and negative binomial (NB) regression to identify potential genomic variables that significantly explain variation in 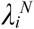 and 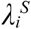, and thus ultimately in the mutation and selection processes from gene-to-gene. The Poisson regression is used to explore counts of rare events (i.e. the observed mutations) that have a fixed probability of being observed, while for a NB regression, the rate of those rare events is itself a random variable that is gamma-distributed. A NB regression incorporates an extra parameter beyond a Poisson rate, known as the dispersion parameter (here denoted by θ), reflecting the amount of underlying variation in the rate of observed mutations from gene-to-gene and governs the “extra” variance of the NB distribution relative to a Poisson distribution with identical mean. If there is no heterogeneity among the rate of observed mutations from gene-to-gene, the dispersion parameter θ goes to zero and we recover a Poisson regression model. Therefore, the Poisson regression model is a special case of the NB regression model, as NB(*λ*_*i*_, θ) reduces to Poisson(*λ*_*i*_) at the limit of θ → 0 (see for instance Zuur, 2009). As a consequence, the Poisson and NB models are “nested” and their relative fit can be compared using a likelihood ratio test when exploring the fit of both types of regression models in this study.

More precisely, we use the models *X*_*i*_ ~ Poisson (*λ*_*i*_) or *X*_*i*_ ~ NB(*λ*_*i*_, θ), fitting the following regression:

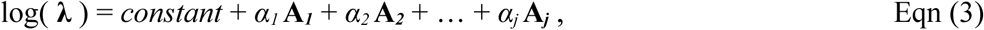

where ***λ*** = (*λ*_*1*_, …, *λ*_*i*_, …, *λ*_*n*_) are the Poisson rates for all *n* genes, ***A***_*l*_, … ***A***_*j*_, are the *j* potential genomic explanatory variables, and *α*_*1*_ … *α*_*j*_, the estimated regression coefficients for those *j* variables. Thus, in the case of the synonymous mutations, *constant* = log(*M*_*0*_ *π*_*0*_ *μ*_*0*_), ***A*_*1*_**, = log(*L*_*i*_) setting *α*_*1*_ = 1. For nonsynonymous mutations, *α*_*2*_ ***A***_*2*_ + … + *α*_*j*_ ***A***_*j*_, = log(*π*_*i*_). Details of the implementation of these models is provided below.

## Methods

### The data

#### Evolution experiment data

We analyzed data obtained from whole genome re-sequencing of forty populations of *S. cerevisiae* adapted in parallel in the lab (Lang et al., 2013). In this experiment, clonal haploid yeast populations were grown in 128 μl of liquid YPD media and transferred every 12 to 24 hours to fresh media for approximately 1,000 generations (see Lang et al., 2011 for more details on the experiment protocol). In our analysis we focus on all detected genic mutations (718 out of the total 1020 in the data set) from forty sequenced populations, i.e. all genic mutations that were able to escape drift and so rise to frequencies of at least approximately 10% in the populations (mutations below this frequency could not reliably be detected, see Lang et al., 2013). The mutations included in the data set consisted of SNPs and small indels. Mutations were grouped by gene across all forty populations, and categorized as synonymous (SYN) or nonsynonymous (NS), i.e. those that do not confer amino acid changes, and those that do, respectively.

#### Comparative genomics data

We used a set of orthologuous gene alignments spanning four distinct yeast species (*S. cerevisiae*, *S. paradoxus*, *S. bayanus*, and *S. mikatae*; available from www.yeastgenome.org/download-data/genomics; Kellis et al., 2003; Cliften et al., 2003) to infer the gene-to-gene heterogeneity of the substitution rates at synonymous sites and nonsynonymous sites, hereafter *dS* and *dN* respectively. To do so, we first realigned the gene sequences using ClustalW (Larkin et al., 2007) on the translated protein sequence data and then applied a number of filters to the data with an aim at removing those gene alignments that might result in inaccurate codon substitution model predictions. We removed alignments for those genes where sequences were not available from all four species, alignments for which at least one sequence had <30% overlap with the one of the other 3 sequences, and alignments for which at least one sequence was <300 bps in length. We then used a maximum likelihood codon based method (CodeML in the PAML software package; Yang, 2007) to infer *dS* and *dN*, for each gene in our data set. We used a codon table model (i.e. seqtype = 1; CodonFreq = 3) with a fixed tree topology (i.e. runmode = 0). A comparison of AICs among alternative codon based models indicated this was the most appropriate model for the data set.

#### Additional genomic data

We included eight additional genomic variables in our analysis that we expected could have the potential to effect the probability of a gene to harbor mutations. Our collection of variables is not meant to be exhaustive, but simply meant to illustrate the potential for additional genomic information to improve our predictions of which genes bear mutations across the genome. For each gene we consider: gene length, % GC content, multi-functionality, degree of protein-protein interaction (PPI), codon adaptation index (CAI), number of domains, level of expression (in the same environment at the evolution experiment), local recombination rate, and essentiality of the gene. We expect some of these variables may capture heterogeneity in the per-gene mutation rates, for example: gene length, which likely captures variation in a gene’s mutational target size, and local recombination rate, which has been shown to be associated with mutability in yeast (Holbeck and Strathern, 1997; Strathern et al., 1995). We expect other variables may capture heterogeneity in selection from gene-to-gene, for example: multi-functionality and PPI, which may characterize aspects of how pleiotropic a given gene is and so the level of evolutionary constraint it is under. We expect still other genomic variables may capture heterogeneity in both mutation and selection. For example, level of expression of a gene may be correlated with gene-to-gene variation in selection as highly expressed genes have been shown to be more highly conserved, both specifically in yeast (Drummond et al., 2005; Pál et al., 2001) and as a more general phenomenon across species (Drummond and Wilke, 2008). On the other hand, level of expression of a gene has also been shown to be positively correlated with mutability (Ness et al., 2015). Descriptions of the variables used in this study and sources from which the data were obtained are provided in Table 1.

**Table 1:** Genomic variables used in this study.

| Variable name | Description | Reference |
| --- | --- | --- |
| *dS* | Number of synonymous substitutions per synonymous site, estimated from gene alignments of *S. cerevisiae*, *S. paradoxus*, *S. bayanus*, and *S. mikatae* (Cliften et al., 2003; Kellis et al., 2003). | Estimated for this study. |
| *dN/dS* | Number of nonsynonymous substitutions per nonsynonymous site, estimated from *S. cerevisiae*, *S. paradoxus*, *S. bayanus*, and *S. mikatae* (Cliften et al., 2003; Kellis et al., 2003). | Estimated for this study. |
| Gene length (*L*) | The number of nucleotides. | (Cherry et al., 2012) |
| % GC content (*GC*) | Percentage of nucleotides in the gene sequence that are either guanine or cytosine. | (Cherry et al., 2012) |
| Multi-functionality (*multifunc*) | Number of different GO slim categories assigned to a gene. | (Cherry et al., 2012) |
| Degree of protein-protein interaction (*PPI*) | The number of physical interactions reported by BioGRID (Stark et al., 2006). | (Koch et al., 2012) |
| Codon adaptation index (*CAI*) | A measure of bias in the usage of synonymous codons, based on a comparison between codon frequencies in the gene and frequencies observed in a set of highly expressed genes (Sharp and Li, 1987). | (Koch et al., 2012) |
| Number of domains (*num.dom*) | The number of regions that Pfam (Punta et al., 2012) has identified as domains in the protein sequence of each gene. | (Koch et al., 2012) |
| Level of expression (*expr*) | A measure of mRNA level for each gene when grown in standard lab conditions. | (Holstege et al., 1998) |
| Local recombination rate (*r*) | Mean recombination rate for a given gene calculated from recombination rate estimate at 0.5 kb intervals using *LDhat* (McVean et al., 2004). | (Illingworth et al., 2013) |
| Essential genes (*essential*) | A true/ false indicator variable denoting whether or not a gene is essential, based on growth assays of deletion strains. | (Winzeler et al., 1999) |

For the purposes of our analysis, we only used mutations in genes for which we were able to obtain a full set of complementary genomic variables (393 of the total 718 genic mutations in Lang et al., 2013). This final data set does not contain any examples of multiple mutations within the same gene on the same genome and so we consider each mutation to be an independent mutational event. A data set integrating the mutation counts originally made available by Lang et al., 2013 (from their Supplementary Table 1) and all the genomic covariates that we aggregated for this study, as well as the gene alignments used for estimating *dN* and *dS* are available on Dryad (doi will be inserted here).

### Regression models

#### Regression models and explanatory variables tested

We used the Poisson and negative binomial regression models described in the “Models” section above to examine how much of the variation in our explanatory variables could account for patterns of variation in mutation counts per gene. We used the ‘glm’ and ‘glm.nb’ functions in R (R Development Core Team, 2014) to implement these models. We fit a series of models to synonymous and nonsynonymous mutation count data separately. To start, we fit the synonymous mutations (model M_S_), testing our assumptions that rate of observed mutations per gene (totaled over all 40 populations in the data set) is proportional to number of nucleotide sites in the gene (*L*_*i*_), and the per nucleotide mutation rate does not vary significantly across the genome - i.e. a model assuming *μ*_*0*_ is a fixed parameter (Poisson regression) fits the data better than a model where *μ*_*0*_ for each gene is drawn from a gamma distribution (NB regression). We also tested for significance of each of the genomic variables included in this study by adding each of them to the Poisson model and checking if the model fit is significantly improved.

After these assumptions were checked, we moved on to fit the nonsynonymous mutation data (M_N_), testing the 11 genomic variables listed in Table 1. We then examined an alternate model (M_N_^PC^), fitting the nonsynonymous mutations using the principal components of the 11 genomic variables in place of the raw variables. The reason we explore this model is that many genomic variables tend to be correlated (for correlations between the particular variables used in this study, see supplementary Table S1), and one approach to reducing potential problems with co-linearity is to transform the raw variables into their principal components and use the resulting uncorrelated composite variables for the regression analysis. We performed a principal component analysis on 11 genomic variables using the ‘prcomp’ function in R to obtain 11 principal components (PCs).

#### Model selection and significance of variables

For each variable and parameter of interest we tested significance by comparing versions of the models with and without that variable or parameter of interest through a likelihood-ratio test (LRT). Significance testing for LRTs was done using permutation tests instead of relying on asymptotic distribution of the LRTs. Permutation was performed by re-sampling the mutation data, re-fitting the models with and without the genomic variable or parameter of interest and then calculating the LRT of those re-fitted models. This procedure allowed us to approximate null distributions and obtaining P-values by calculating the frequency of permutations that resulted in a likelihood-ratio greater than or equal to the true observed value. Variables found to significantly improve model fit were retained in the final “best” model. We choose to test significance using permutations given that asymptotic results on the distribution of the likelihood ratio test may break down as the reduced model – the Poisson regression – lies at the boundaries of the parameter space for θ, included in the NB regression (see for instance Self and Liang, 1987). In practice, 1000 permutations were used to approximate the null and obtain p-values on each variable (more permutations might be required if needed to approximate p-values that are much smaller than 10^-3).

The two nonsynonymous mutation models M_N_ and M_N_^PC^ were compared with each other using Akaike information criterion (AIC; Akaike, 1973), and the proportion of variation explained (pseudo-R^2^) was estimated as the R^2^ obtained from a linear regression (using ‘lm’ in R) between the observed and predicted mutation counts for a given model. Note that this statistic is not used for any formal goodness-of-fit but as an illustrative way to report how much of the whole variation is accounted for by any model we fit to the mutation count data.

#### Predicting the degree of parallelism

We used the resulting best-fit parameters and distributions from the regression models to simulate mutations for 40 populations and calculated the mean proportion of shared genes bearing mutations for all pairwise combinations of those 40 populations using the Jaccard Index, 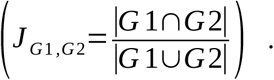 This measure of parallelism has been used in a number of previous empirical comparisons (Bailey et al., 2017; Wong et al., 2012). We compared the simulated distribution of J values to the distribution calculated from the real data set in order to assess the accuracy of the regression models in predicting the degree of parallel evolution in this system. This can be used as a predictive check for our model.

All statistical analyses were implemented in R (R Development Core Team, 2014) and an example script for implementing our model framework and hypothesis testing is available on Dryad (doi will be inserted here).

## Results

### The data

#### Mutation counts data

We used experimental data comprising all mutations detected at a frequency over 10% in the forty evolved *S. cerevisiae* populations described in Lang et al., 2013. After removing those genes for which we had incomplete or unreliable data (see Methods), we were left with 2891 genes out of a total of 6603. The filtered data set contained 357 nonsynonymous mutations distributed across 267 genes, and 58 synonymous mutations distributed across 57 genes. The genes removed by our filtering rules had disproportionately more mutations compared to those genes that were retained in the data set (*X*^2^ = 50.57, df = 1, P < 0.001). This is not unexpected as highly divergent genes are more likely to be filtered out due to alignment issues, and it is not surprising that highly divergent genes would tend see more mutations than average, whether it be as a result of mutation and / or selection mechanisms. This bias in the filtering means that our results are likely conservative in terms of detecting significant relationships between long-term (from comparative genomics data) and short-term (from experimental evolution data) measures of divergence.

#### Genomic variables

We used codon substitution models comparing four yeast species to estimate *dS* and *dN/dS* for each gene. Estimates for *dS* ranged widely, from 0.21 to 68.7, however the vast majority of *dS* estimates (~95%) were less than 4. Estimates for *dN/dS* ranged from 0.00010 to 0.43, and these values are weakly negatively correlated with *dS* (r = -0.043, P = 0.021). We collated and/ or calculated nine other genomic variables with the potential to effect the mutation and selection processes in this system and estimated correlation coefficients between all pairs of explanatory variables used in this study (Table S1). While the correlations between these variables tend to be quite weak, many are, in fact, significant due to the large number of observations in the data set.

### Mutation counts analysis

#### >Synonymous mutations

We used regression models to test our assumption that gene-level mutation rate can be adequately described as simply being directly proportional to gene length. Restricting the data to the synonymous mutations, we compared Poisson regression models with and without gene length included as an explanatory variable (M_S_0.P: λ_S_ = *constant* and M_S_1.P: *λ*_S_ = *constant*\*(*L*_*i*_)^α1^, respectively), and a Poisson regression model where rate is restricted to be directly proportional to gene length (i.e. M_S_2.P: *λ*_S_ = *constant*\**L*_*i*_). We also compared with negative binomial versions of these model to look for the possibility of additional unexplained variation in the rate *λ*. A summary of the results of these comparisons is shown in Table 2. Model M_S_2.P was the best model according to a comparison of AICs. The fits of these models to the distribution of synonymous mutation counts per gene are visualized in Fig. 2A. We also compared a series of Poisson models, each containing one of the genomic variables included in this study (see supplementary information Table S2). Two variables do significantly improve model fit – dN/dS and CAI, but very modestly, helping to explain only 0.24 % and 0.11 % of the total variance respectively. Taken together, these tests suggest that the assumptions of neutral selection at synonymous sites and a constant mutation rate across nucleotides are reasonable simplifications for these data.

**Table 2:** ‘M_S_’ models testing assumptions with the synonymous mutation data. Log-likelihoods, and AIC values are provided. The best model as determined by the lowest AIC with the fewest parameters is highlighted in grey.

| Model |  | log-lik. | No. param. | AIC |
| --- | --- | --- | --- | --- |
| M<sub>S</sub>0.P: | Pois( $\lambda_S = \text{constant}$ ) | -283.0 | 1 | 568.2 |
| M<sub>S</sub>0.NB: | NB( $\lambda_S = \text{constant}, \theta_S$ ) | -283.0 | 2 | 569.9 |
| M<sub>S</sub>1.P: | Pois( $\lambda_S = \text{constant} * L_i^{\alpha 1}$ ) | -273.9 | 2 | 551.8 |
| M<sub>S</sub>1.NB: | NB( $\lambda_S = \text{constant} * L_i^{\alpha 1}, \theta_S$ ) | -273.9 | 3 | 553.8 |
| M<sub>S</sub>2.P: | Pois( $\lambda_S = \text{constant} * L_i$ ) | -274.0 | 1 | 549.9 |
| M<sub>S</sub>2.NB: | NB( $\lambda_S = \text{constant} * L_i, \theta_S$ ) | -274.0 | 2 | 551.9 |

**Figure 2:**
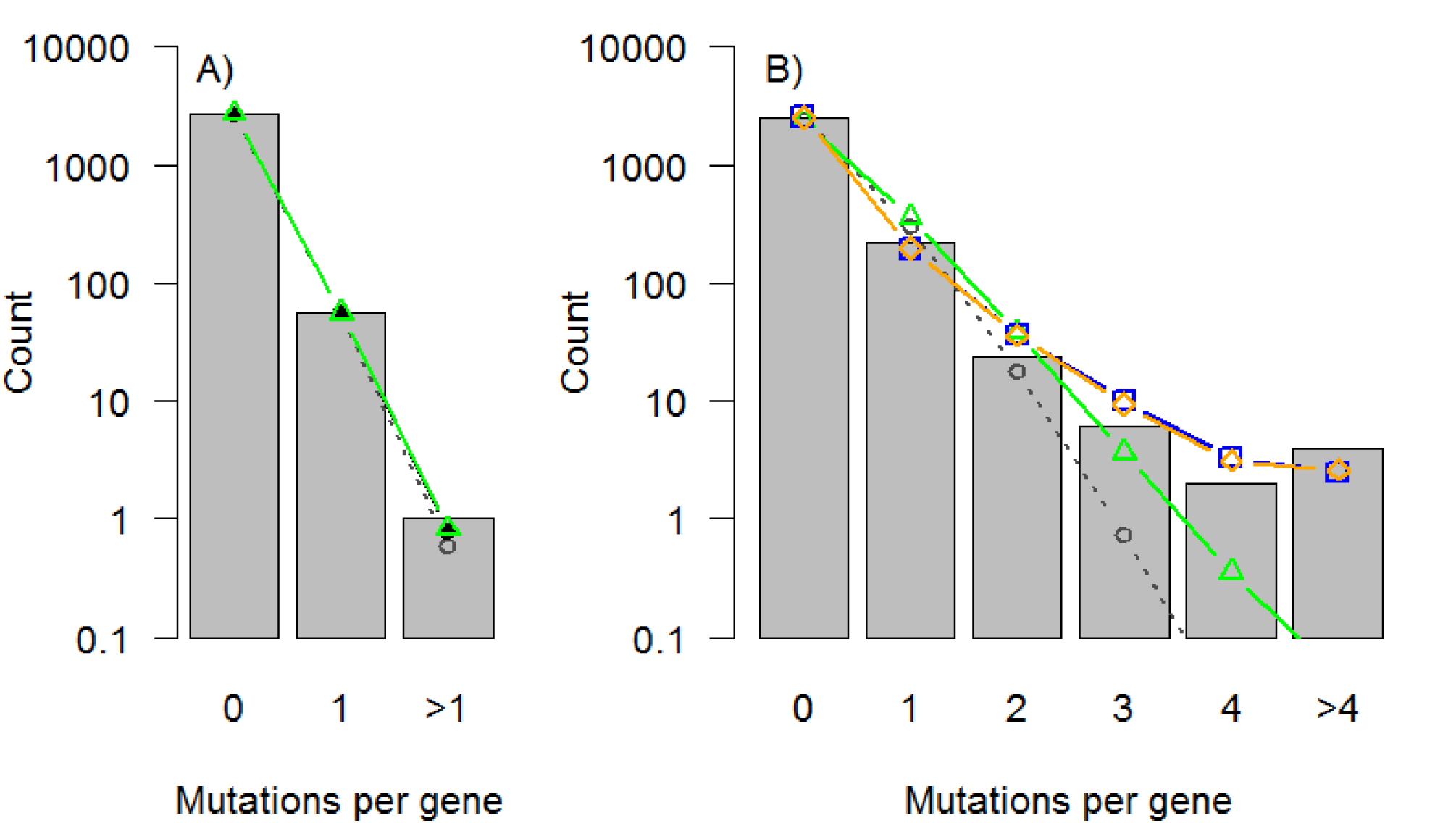
Distribution of A) synonymous and B) nonsynonymous mutations per gene (totaled over all 40 populations in the data set) and predicted model distributions from M0.P (grey circles), M1.P (black points), M2.P (green triangles), and M_N_.NB (blue squares), and M_N_.NB_PC_ (orange diamonds).

#### Nonsynonymous mutations

We fit regression models to the nonsynonymous mutation data, including eleven genomic variables, trying to identify which of those variables could significantly explain variation in the number of observed mutations per gene (totaled over all 40 populations in the data set). We found that gene length (*L*), number of domains in the encoded protein (*num.dom*), and recombination rate (*r*) were significant in our model (see model M_N_.NB in Table 3).

**Table 3:**
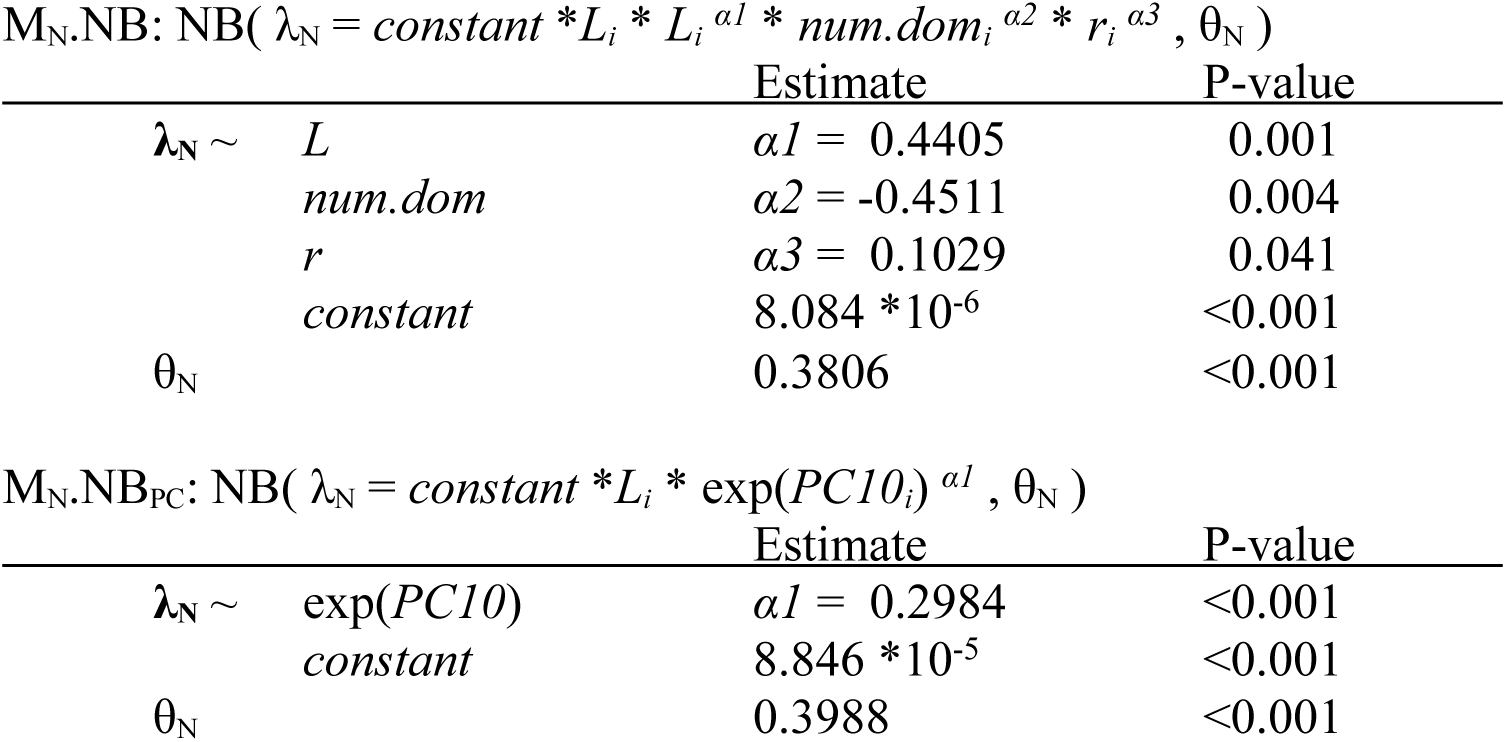
‘M_N_’ models parameter estimates (*constant, α1, α2, etc*) and P-values for those estimates. Only those variables that significantly improved model fit are included.

| M<sub>N</sub>.NB: NB( $\lambda_N = constant * L_i * L_i^{ \alpha 1} * num.dom_i^{ \alpha 2} * r_i^{ \alpha 3} , \theta_N$ ) | | | |
| --- | --- | --- | --- |
|  |  | Estimate | P-value |
| $\lambda_N \sim$ | $L$ | $\alpha 1 = 0.4405$ | 0.001 |
| | $num.dom$ | $\alpha 2 = -0.4511$ | 0.004 |
| | $r$ | $\alpha 3 = 0.1029$ | 0.041 |
| | $constant$ | $8.084 * 10^{-6}$ | <0.001 |
| $\theta_N$ | | 0.3806 | <0.001 |

| M<sub>N</sub>.NB<sub>PC</sub>: NB( $\lambda_N = constant * L_i * \exp(PC10_i)^{ \alpha 1} , \theta_N$ ) | | | |
| --- | --- | --- | --- |
|  |  | Estimate | P-value |
| $\lambda_N \sim$ | $\exp(PC10)$ | $\alpha 1 = 0.2984$ | <0.001 |
| | $constant$ | $8.846 * 10^{-5}$ | <0.001 |
| $\theta_N$ | | 0.3988 | <0.001 |

When we fit regression models using the principal components of the genomic variables in place of the raw variables, we found that only a single principal component, PC10, was significant in the model (see model M_N_.NB_PC_ in Table 3). PC10 is fairly evenly loaded with a number of genomic variables (see Fig. 3), however the three significant genomic variables from M_N_.NB (*L*, *num.dom*, and *r*) are among the variables more heavily loaded on PC10, so the two models seem to be roughly in agreement. Further, these models both explain about 16% of the variance (as calculated from pseudo-r^2^ estimates, see methods). A comparison of Poisson and negative binomial regression models, as well as models including the raw genomic variables versus the transformed principal component variables, suggests that the best model for these nonsynonymous mutation count data is a negative binomial regression using the raw genomic variables (see AIC values in Table 4). The fits of these models to the distribution of nonsynonymous mutation counts per gene are visualized in Fig. 2B.

**Figure 3:**
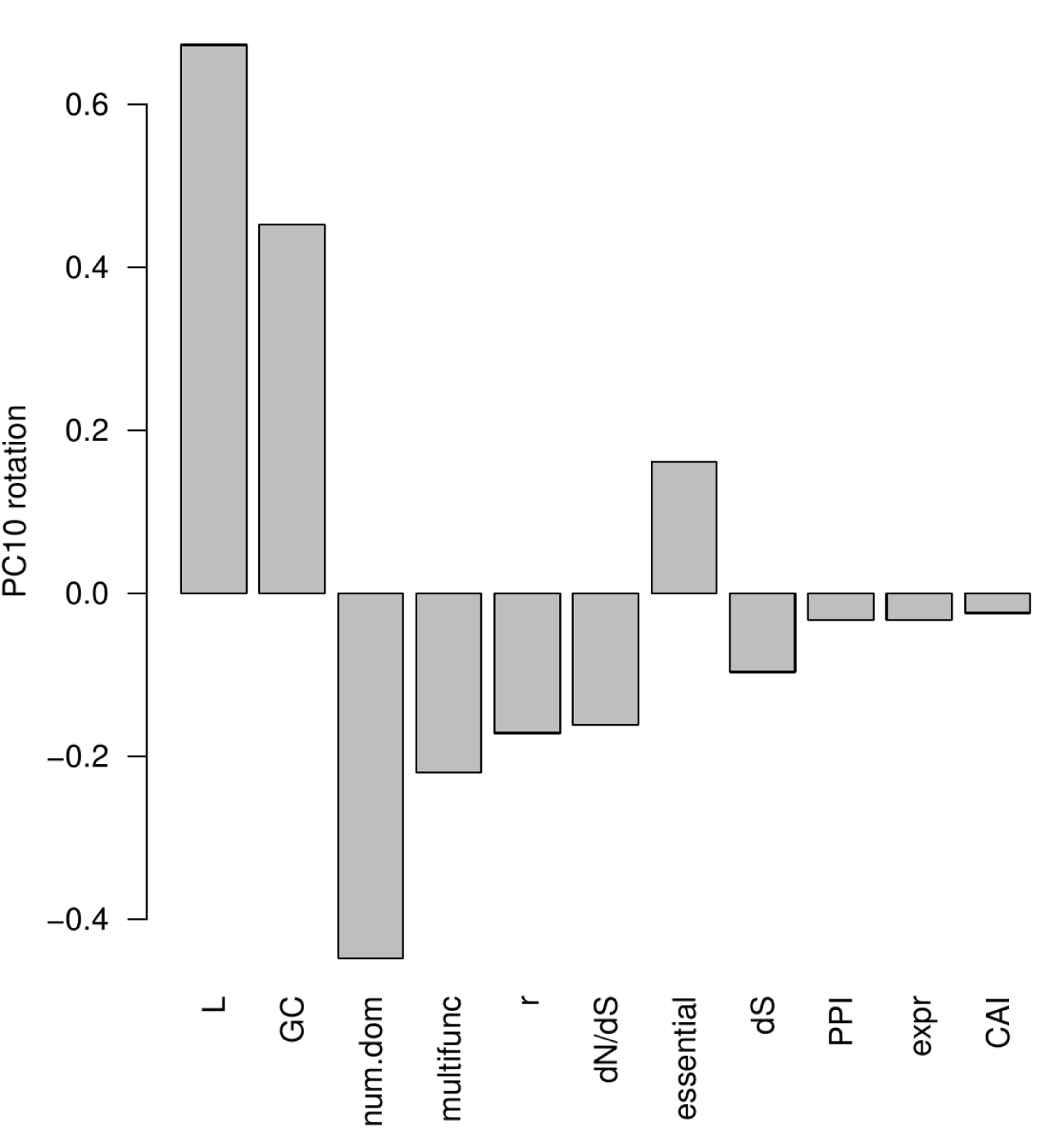
Loadings of the 11 genomic variables on PC10 – the only principal component that significantly explains variation in nonsynonymous mutation counts. Genomic variables are ordered from largest to smallest in terms of the absolute value of their loading.

**Table 4:** Log-likelihoods, and AIC values for the ‘M_N_’ models. The best model as determined by the lowest AIC with the fewest parameters is highlighted in grey.

| Model | log-lik. | No. param. | AIC |
| --- | --- | --- | --- |
| M<sub>N</sub>0.P: | -1159.9 | 1 | 2321.7 |
| M<sub>N</sub>0.NB: | -1022.5 | 2 | 2048.9 |
| M<sub>N</sub>2.P: | -1050.3 | 1 | 2102.7 |
| M<sub>N</sub>2.NB: | -956.6 | 2 | 1917.3 |
| M<sub>N</sub>.P | -1021.4 | 4 | 2050.8 |
| M<sub>N</sub>.P<sub>PC</sub> | -1013.0 | 2 | 2030.1 |
| M<sub>N</sub>.NB | -944.8 | 5 | 1899.7 |
| M<sub>N</sub>.NB<sub>PC</sub> | -947.9 | 3 | 1901.8 |

#### Predicted parallelism

Using the predicted rates and distributions from the best-fit regression models for both the synonymous and nonsynonymous mutations (M_S_2 and M_N_.NB, respectively), we simulated mutations for a set of 40 populations and calculated the Jaccard index (J) as a measure of gene-level parallelism between all pairs of those simulated populations. We performed the same calculations with the 40 populations from the real data. Fig. 4 shows a comparison of those J values from the simulated and real data. While the simulated data from our model does quite well at capturing the range of J values, it does not do a great job of capturing the shape of the distribution.

**Figure 4:**
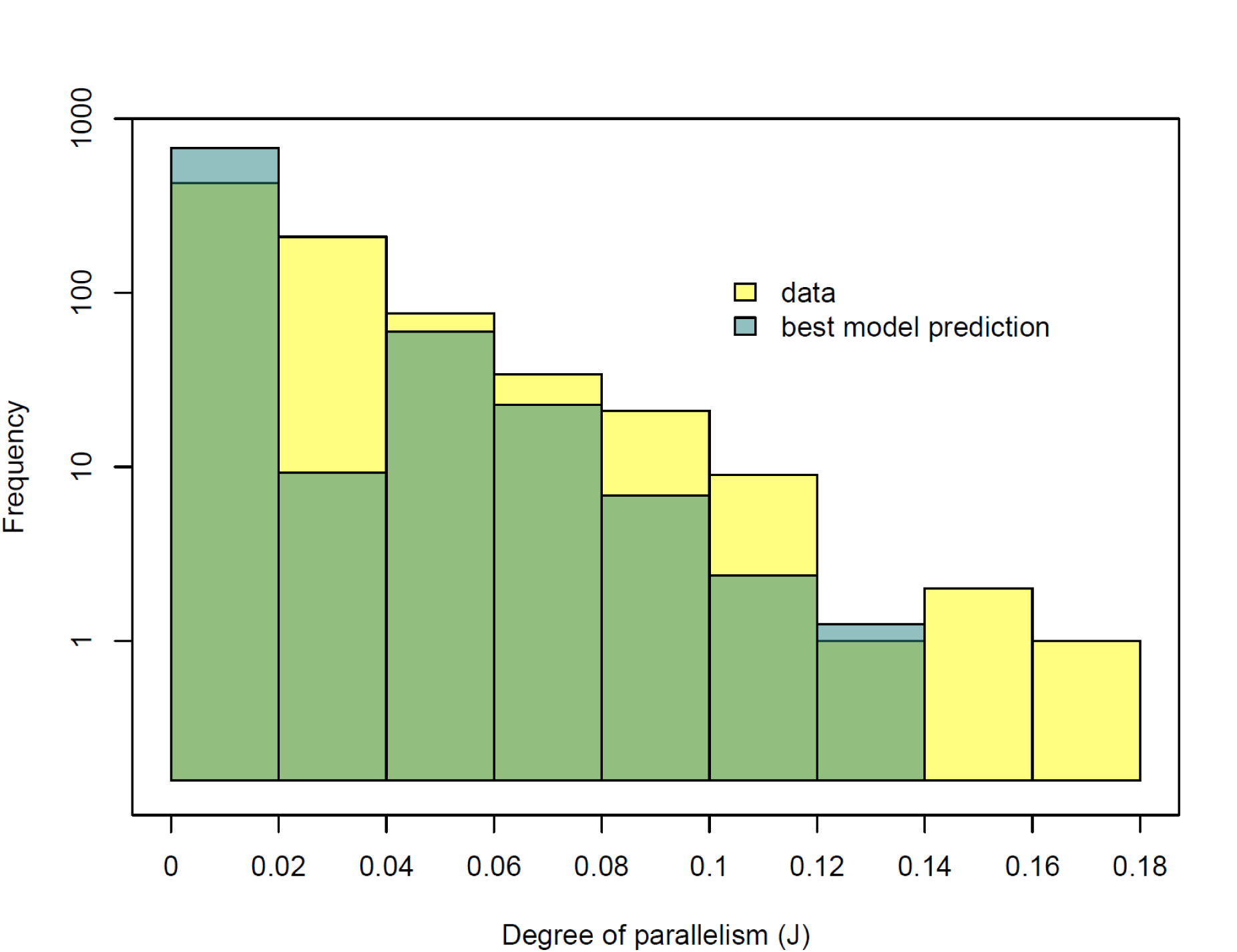
Distribution of the degree of parallelism (estimated as the pairwise Index, J) from the real data and simulated data from the best-fit models.

## Discussion

Here we present a modeling framework to infer what genomic variables may underlie gene to gene variation in mutation rate and intensity of selection. We use these models to provide evidence that parallel evolution at both nonsynonymous and synonymous sites is driven by non trivial amounts of gene-to-gene heterogeneity in the mutation and selection processes. Using our modeling approach, we identified a number of genomic variables that can significantly improve models predicting the distribution of mutations observed across genes in experimentally evolved populations of *S. cerevisiae* (Lang et al., 2013). We are also able to classify genomic variables into those that have affected mutation counts 1) through their effect on the mutation rate (variables that significantly predict synonymous mutations), and/ or 2) through their effect on the probability of a mutation being either observed/ lost due to selection (variables that significantly predict nonsynonymous mutations). Out of all the variables tested, we found that *gene length* explained the most variation in both synonymous and nonsynonymous mutation counts per gene – plainly speaking, longer genes accumulate more mutations. However, *number of domains* and *recombination* also had significant effects. Below we discuss in detail these genomic variables and their potential contributions to the probability of parallel evolution via the processes of mutation and selection.

### Longer genes harbor more mutations

By far, the variable having the largest effect on variation in the number of synonymous and nonsynonymous mutations observed was *gene length*. More specifically, gene length positively affected the rate of mutation at the gene-level, meaning genes comprising more nucleotides were more likely to harbor mutations. This result is not surprising and is in agreement with recent analysis of synonymous mutation counts from Lenski’s long term evolution experiment with *E. coli* (Maddamsetti et al., 2015).

### Long-term divergence does not predict short-term mutation counts

Our model for synonymous mutation counts suggests that divergence estimates from long-term evolutionary comparisons at the species level do not provide insight into expected mutation counts on the shorter time scale of evolution in the lab, also in agreement with recent analysis of *E. coli* data (Maddamsetti et al., 2015). Maddamsetti et al found that their proxy for long-term per gene mutation rate, θ_s_ (a measure of within-species nucleotide diversity), did not explain gene-to-gene variation in synonymous mutation counts in their data. The authors argued that horizontal gene transfer (HGT) is therefore likely a more important process driving gene-to-gene variation in long-term divergence between naturally occurring *E. coli* strains, and since HGT did not occur in their evolution experiment, it is not surprising that the experiment’s synonymous mutation counts did not correlate with θ_s_. However, rates of HGT tend to be higher in bacteria, and in particular *E. coli*, as compared to yeast and other eukaryotes (e.g. Boto, 2010). Furthermore, a recent mutation accumulation experiment with the eukaryote *Chlamydomonas reinhardtii* showed a positive correlation between a proxy for long-term mutation rate (θ_s_) and per site mutability (Ness et al., 2015). Thus, it is somewhat surprising that we do not see a significant relationship between *dS* and *dN/dS* and counts of synonymous and nonsynonymous mutations respectively in our examination of the *S. cerevisiae* data used in this study. One possibility might also be that *dS* and *dN*/*dS* are noisy to estimate at the gene level and that tends to downplay their predictive power in our analysis of counts in an evolve and re-sequence experiment.

### Nonsynonymous mutation counts show evidence of selection heterogeneity

As expected (Lenormand et al., 2016), we see strong evidence that the distribution of nonsynonymous mutations across the genome is driven in part by gene-to-gene heterogeneity in selection. Of those genomic variables tested, we found three that were significant predictors of nonsynonymous mutation counts, suggesting that those variables may drive or are correlated with processes that modulate the intensity of selection across genes. The significant variables were *gene length*, *recombination rate*, and *number of protein domains*.

We found that *gene length* predicts nonsynonymous mutation count via selection, over and above its effects on per gene mutation rate – as estimated from models aimed at explaining the synonymous mutation count only. While one might not expect *gene length* to have direct effects on selection, we suggest that *gene length* may show a significant effect here because it is correlated with other attributes of the genome that could have important effects on selection, for example *gene expression levels* and *multifunctionality*. Because of these correlations, it could be that *gene length* acts as a kind of summary variable for these covariates and other unidentified factors we have not captured in these models. In fact, it is almost certainly the case that to some degree all the significant variables in our model summarize variation from additional unknown factors we have not included in our data set.

In contrast to the positive relationship between *gene length* and number of nonsynonymous mutations, we also found that the *number of protein domains* that a gene codes for (a variable that is positively correlated with gene length; Table S1) actually negatively predicts the number of nonsynonymous mutations. In other words, the more domains in the encoded protein of a gene, the fewer mutations that gene is expected to incur in the course of the yeast evolution experiment analyzed here. The mechanism behind this effect is not clear, but certainly protein structure has previously been reported to have significant impacts on evolutionary rates in yeast (Bloom et al., 2006) and one can also posit that genes encoding proteins with multiple domains and thereby involved in more numerous interactions are – all else being equal – more severely constrained by purifying selection. It is interesting that this effect can be observed in the course of relatively short time span (relative to between species divergence times) through the relative paucity of nonsynonymous mutations in these genes.

Our analysis also showed that *recombination rate* is a significant predictor of the number of nonsynonymous mutations observed in a given gene in these data. Genes with higher recombination rates are more likely to bear nonsynonymous mutations. We expect recombination rate to be correlated with mutation, as previous studies in yeast have shown that recombinational repair of double strand breaks substantially increases the frequency of point mutations in nearby intervals (e.g. Holbeck and Strathern, 1997; Strathern et al., 1995). However, it is not clear how high recombination rates might drive, or be correlated with other processes that drive, selection – as our models suggest is the case for this data set. Another non-exclusive possibility might be the fact that biased gene conversion might vary from gene to gene and also – like selection – affect the probability of detecting variants in evolve and re-sequence experiments. However, if this was the case, we might expect GC content to be a significant predictor of both synonymous and nonsynonymous mutations in our models, and it is not (S: P = 0.125, NS: P = 0.221). We might also expect to see a bias towards GC in the observed mutations, however this was not the case – only 33% of the SNPs in this data set were a change to G or C.

### Factors driving mutation and selection are complex

It is difficult to obtain any additional insights from our models that include principal components of the genomic covariate data, however there is at least some level of agreement between those variables that are significant (i.e. length, recombination, and number of domains) and ones that are heavily weighted in PC10 – the principal component that was found to be significant (see Fig. 3). The local properties of the genome do appear to drive some heterogeneity in the selection processes, and in turn, shape the patterns of parallel evolution, however individual effects that can be ascribed to individual variables are not easy to parse out.

We want to stress that while we were able to identify a number of factors affecting the count of mutations observed in this evolution experiment data set, the total explained variance is still low: 1 % and 16 % in the synonymous and nonsynonymous models respectively (calculated from pseudo-r^2^ estimates of the “best” models, see methods). While the models do capture the general distribution of mutation counts (Fig. 2) and so the degree of parallel evolution, accurately predicting on which genes those mutations will fall is still very difficult. This is not surprising given the amount of stochasticity involved in both the origin of new mutations and their evolutionary fate through drift and selection. A clearer picture might emerge when using our modeling approach in a meta-analysis approach where several evolve and re-sequence experiments are considered together (see Bailey et al., 2017 for a similar approach on summary statistics of the amount of parallel evolution at the gene level across a wide range of experimental studies in yeast and bacteria).

While we do find a number of genomic variables that significantly affect the distribution of mutations across the genome, it is noteworthy that these models are still unable to capture the more extreme patterns of parallel evolution observed in this data set. For example, one gene (IRA1) saw mutations in over 50% of the populations sequenced in this experimental data set (discussed in more detail in Lang et al., 2013). Such a mutation count is completely out of the range of likely outcomes predicted by our models. Some of this discrepancy may be because of the simplifying assumptions made about the process of selection. Our framework models the process of mutation and its heterogeneity but while we account for the fact that newly arising mutations may have different probabilities of reaching an observable frequency, the modeling of that process could be made more precise by incorporating an explicit underlying distribution of fitness effects of new mutations at each gene. Incorporating a selection process that allows for different amounts of both positive and negative selection, as well as further details about the selection pressures in the particular environment of interest – something we do not consider at all in this study – would likely improve prediction for some of these more extreme events.

It is also important to note that the methods used in this study are focused on parallelism in SNPs and small indels. While this focus is appropriate for the data set used here, it may not be appropriate for other systems. For example, in an experimental evolution study with E. coli, (Tenaillon et al., 2012) saw that much of the parallelism seen between populations was the result of IS elements and large scale duplication and deletions. It may be important to try to account for this more diverse range of mutational event types when trying to identify the drivers of parallel evolution other systems.

Can we use this modeling framework to predict parallel evolution? To some degree, yes – the measures of parallel evolution between populations simulated using our model predictions span a similar range to those calculated from the real population data (see Fig. 4). However, while this congruence suggests we are on the right track, the shapes of the real and simulated distributions are still quite different. For example, there is, on average, more parallelism between the real populations compared to populations simulated from our models, and in particular, there seems to be a substantial discrepancy between the number of real population-pairs and simulated population-pairs that have a low level of parallelism (i.e. note the difference between real and simulated when J ranges from 0.02 to 0.04 in Fig. 4). This is further evidence suggesting that although our current models may be useful to some extent, we are still missing some important factors driving heterogeneity in mutation and selection across these genomes.

### Advantages of this regression framework

Relying on the assumption that synonymous mutations are selectively neutral (which does appear to be appropriate for these data), the regression models we use in this study allow us to distinguish between genomic variables influencing the observed distribution of mutations across a genome through their potential effects on both gene-to-gene heterogeneity in mutation rate and gene-to-gene heterogeneity in selection. The great advantage of this is that it allows us to begin to break down the importance of these two processes in shaping patterns of parallel evolution we see, and move closer to the goal of predicting which genes will be involved in evolution when organisms adapt to new environments. It will be interesting to apply this model framework to other data sets of this type, as they become available, to see how general these patterns are across different organisms and selection environments (Bailey and Bataillon, 2016).

## Acknowledgments

We thank three PCI Evolutionary Biology recommenders for their constructive comments. This work was supported by the European Research Council under the European Union’s Seventh Framework Program [FP7/20072013, ERC grant number 311341 to T.B.].

